# Long-read DNA sequencing reveals global hypomethylation and focal hypermethylation in *SDHx*-deficient pheochromocytoma and paraganglioma

**DOI:** 10.64898/2026.11.04.763174

**Authors:** Zhuwei Xu, Hemant K. Prajapati, Christopher T. Coey, Peter R. Eriksson, Igor Hartmann, Hussam Alkaissi, Karel Pacak, David J. Clark

## Abstract

Pheochromocytomas and paragangliomas (PPGL) are highly heritable neuroendocrine tumors of the autonomic ganglia. Genomic DNA in most cancer cells is hypomethylated. However, DNA in PPGLs secondary to mutations in succinate dehydrogenase complex (*SDHx*) genes is reportedly hypermethylated due to succinate-mediated inhibition of DNA demethylases. We used nanopore long-read sequencing to characterize the methylome of tumor and white blood cell DNA from *SDHx* and non-*SDHx* PPGL patients, alongside normal adrenal medulla. We find that both *SDHx* and non-*SDHx* PPGLs are globally hypomethylated, particularly at centromeres. In normal adrenal medulla, CpG islands marked by H3K4 methylation are hypomethylated, whereas CpG islands lacking H3K4 methylation are hypermethylated. In *SDHx*-related PPGL, some H3K4me-marked CpG islands are hypermethylated, but most are not. We propose that DNA methylation in *SDHx*-related PPGL reflects the global hypomethylation generally observed in tumors, partly offset by succinate-mediated inhibition of TET DNA demethylases. Opposing effects of succinate may alter the balance between DNA methylation (indirect inhibition of the DNMT3A/B DNA methylases via inhibition of H3K4me demethylases) and DNA demethylation (inhibition of TET DNA demethylases) at active CpG islands.

## Introduction

Pheochromocytomas are neuroendocrine tumors derived from chromaffin cells in the adrenal medulla. Paragangliomas derive from sympathetic or parasympathetic ganglia external to the adrenal glands and are more likely to metastasize ^1, 2^. Pheochromocytomas and paragangliomas (PPGL) are relatively rare, affecting around 6 per million individuals, but are highly heritable tumors^3, 4^.

PPGLs are associated with mutations in a number of genes, which have been divided into three transcriptomic clusters ^5^: the pseudohypoxia cluster (Krebs cycle genes, such as *SDHx* and *FH,* as well as hypoxia-related genes, such as *VHL, EGLN1/2* and *EPAS1*), the kinase signaling cluster (*RET, HRAS, NF1, TMEM127, FGFR1*) and the Wnt-altered cluster (*MAML3* fusions, *CSDE1* mutations). Germline heterozygous pathogenic variants in succinate dehydrogenase complex genes (*SDHA, SDHB, SDHC, SDHD, SDHAF2*, collectively referred to as *SDHx*) show incomplete penetrance, where some individuals are silent carriers and others develop tumors characterized by loss of heterozygosity. Examples are PPGL, renal cell carcinoma (RCC), gastrointestinal stromal tumors (GIST) ^6, 7^ and pituitary tumors ^3, 8^. The highest penetrance is seen in *SDHB* families, reaching up to 40% ^9^. Individuals with *SDHB* mutations are also at highest risk of metastases ^3, 9, 10, 11, 12^. Metastasis appears to require a secondary mutation in the primary tumor, including *ATRX/TERT* alterations ^5, 13^.

Succinate dehydrogenase (SDH) is a hetero-tetramer in which the SDHB subunit connects the SDHC and SDHD subunits in the mitochondrial inner membrane to the catalytic SDHA subunit. *SDHx* mutations that reduce SDH enzyme activity result in succinate accumulation ^14, 15, 16^. Excess succinate is transported from the mitochondria to the cytoplasm and the nucleus ^17, 18^, where it competitively inhibits 2-oxoglutarate-dependent dioxygenases such as prolyl hydroxylase ^17, 18^, Jumonji C (JmjC) histone demethylases ^19, 20, 21^ and ten-eleven translocation (TET) DNA demethylases ^20, 22^.

The DNA in most tumors is hypomethylated relative to DNA in normal cells, including non-*SDHx* PPGL tumors ^5, 23^, consistent with inappropriate derepression of many genes in tumors. General hypomethylation stems from lack of maintenance DNA methylation by DNMT1 through passive dilution of existing methylation during DNA replication ^24, 25^, and increased active demethylation by TET demethylases ^26^. In contrast, *SDHx*-related PPGL and GIST tumor DNA is reportedly hypermethylated, particularly at CpG islands ^27, 28, 29^. This hypermethylation is attributed to succinate inhibition of the TET DNA demethylases, resulting in slower global demethylation rates.

Various methods have been used to measure DNA methylation in *SDHx* tumors. The predicted loss of 5-hydroxymethylcytosine (5hmC) in DNA due to succinate inhibition of TET DNA demethylases provides the basis for a sensitive immunohistochemical assay for global DNA demethylation ^30^ by comparison with neighboring cells ^20, 31^. Microarray analysis ^5, 7, 28^ and sequencing of hybrid-selected bisulphite-treated DNA fragments ^27^ have both been used to map DNA methylation in specific genomic regions, such as CpG islands. However, these studies do not provide methylation data for the entire genome. Here, we used nanopore long-read sequencing combined with the Telomere-to-Telomere human genome assembly ^32^ to obtain complete genome sequences with DNA methylation profiles ^33, 34^. We compared *SDHx* tumors, non-*SDHx* tumors and the matched blood samples from these patients with normal adrenal medulla. Although we confirm that some CpG islands are hypermethylated, we find that the *SDHx* genome is globally hypomethylated, and therefore similar to other tumors in this regard. Our observations are inconsistent with the proposal that succinate inhibits TET DNA demethylases, resulting in global hypermethylation.

## Results

### *SDHx*-related PPGL tumor DNA is globally hypomethylated

We performed nanopore long-read whole-genome sequencing on matched tumor and white blood cells from ten patients: five with *SDHx* mutations ("SDH01-05") and five with non-*SDHx* mutations ("nonSDH01-05"). We also sequenced adrenal medulla DNA from five healthy donors as controls ("Normal01-05") (Supplementary Table 1; Supplementary Fig. 1). *SDHx* tumors exhibit lower average methylation compared to normal adrenal medulla (Fig. 1a; Supplementary Table 1). Mean methylation levels in *SDHx* tumors range from 59.0% to 66.3%, compared to 66.1% to 69.8% in normal adrenal medulla. In contrast, the matched white blood cell DNA samples are much more methylated (77.5% to 79.6%). On average, *SDHx* tumor DNA is hypomethylated across all autosomes (Fig. 1b), with a mean difference in chromosomal methylation level of 4.2% (standard deviation (SD) = 1.7%; n = 22 autosomes) relative to normal adrenal medulla, indicating a global effect. We conclude that *SDHx* tumor DNA is hypomethylated relative to blood from the same patient and relative to normal adrenal medulla.

**Figure 1.**
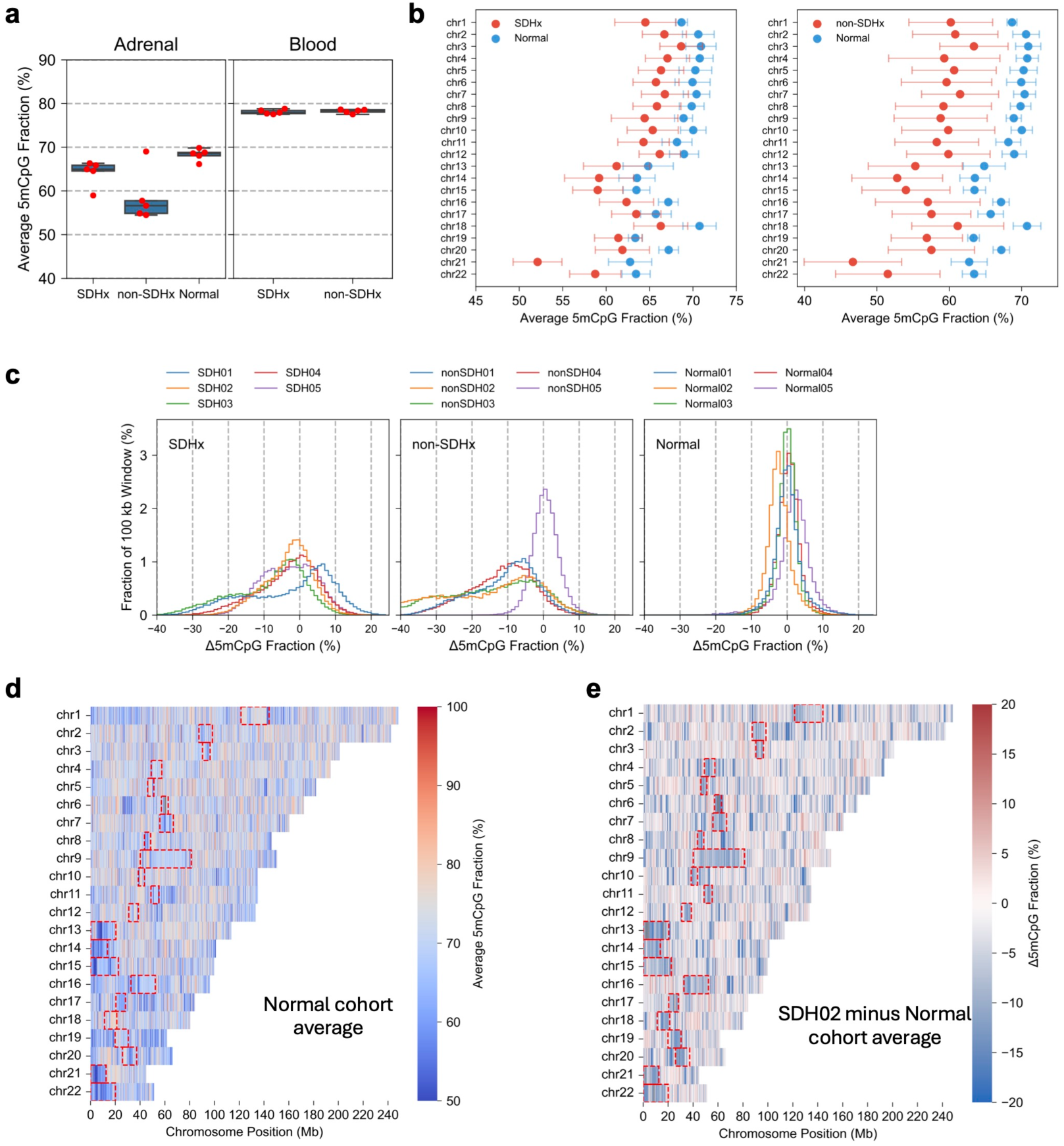
*SDHx*-related PPGL tumor DNA is globally hypomethylated. (**a**) Average CpG methylation (autosomes only). Box boundaries show the 25th and 75th percentiles, the central line indicates the median, and whiskers extend to 1.5x the interquartile range or the most extreme data point within that range. (**b**) Point-range plot comparing mean methylation levels in each autosome in normal adrenal medulla with *SDHx* tumors (left panel) or with non-*SDHx* tumors (right panel). Points indicate the mean methylation of 5 samples per group; error bars indicate standard deviation. (**c**) Genomic distribution of CpG methylation in autosomal 100-kb windows relative to the mean methylation of the normal adrenal medulla cohort for the same window. Histograms for normal adrenal medulla, *SDHx* tumors and non-*SDHx* tumors. (**d**) Genomic methylation heatmap for the normal adrenal medulla cohort. Average methylation in 100 kb windows for the 5 normal samples. Red boxes: centromeres. (**e**) Methylation difference heatmap for the SDH02 tumor. Average methylation in 100 kb windows (SDH02 minus mean normal cohort). Red indicates hypermethylation; blue indicates hypomethylation relative to normal adrenal medulla. Red boxes: centromeres.

Non-*SDHx* tumors are more variable than *SDHx* tumors (54.5% to 69.0% mean methylation). One of the five non-*SDHx* tumors does not show the global hypomethylation observed in most tumors (nonSDH05); the other four tumors are hypomethylated (Fig. 1a; Supplementary Table 1). The wider range in autosomal methylation among non-*SDHx* tumors is clear in the point-range plot showing the average methylation for all five tumors (Fig. 1b). This non-*SDHx* tumor heterogeneity presumably reflects their diverse genetic drivers.

To examine genome-wide methylation changes in detail, we partitioned the 22 autosomes into 100-kb windows and calculated the average methylation for each window. Then we calculated the mean of the normal adrenal medulla cohort for each window and subtracted it from the value for the same window in each sample to obtain a difference plot (Fig. 1c). The five normal adrenal medulla samples exhibit highly concordant methylation patterns ("Normal" panel; mean difference

= -2.2% to 1.9%; SD = 3.0% to 5.1%), with most windows differing by < 10% from the cohort mean (Fig. 1c). Most tumor samples, however, exhibit a wide range of methylation levels relative to the normal adrenal medulla mean ("*SDHx*" and "non-*SDHx*" panels; Fig. 1c). Four of the five *SDHx* tumors are predominantly hypomethylated, whereas SDH01 displays both hyper- and hypo-methylated 100-kb windows. Among non-*SDHx* tumors, four are extensively hypomethylated, whereas nonSDH05 has methylation patterns similar to normal adrenal medulla (Fig. 1c).

Hypomethylated and hypermethylated 100-kb windows are distributed throughout every autosome in the normal adrenal medulla cohort (Fig. 1d). The heatmap shows the mean methylation of each window in each autosome. Many of the hypomethylated windows are located in the centromeres, but not all. Difference heatmaps for each tumor were obtained by calculating the difference between the tumor methylation and the normal adrenal medulla cohort mean methylation for each 100-kb window (Supplementary Fig. 2). SDH02 is shown as an example (Fig. 1e). Hypomethylated windows are predominant in all the tumor samples, although SDH01 shows a larger fraction of hypermethylated windows than the others (Supplementary Fig. 2; Fig. 1c). We conclude that *SDHx* tumor DNA is mostly hypomethylated at the global level.

### Global hypomethylation patterns overlap in PPGL tumors

We compared DNA methylation patterns in 100-kb windows in the *SDHx* and non-*SDHx* tumors relative to normal adrenal medulla (Fig. 2). A beta-binomial statistical framework was used to account for biological variability in the normal cohort. The normal adrenal medulla samples showed low over-dispersion (median intraclass correlation ρ = 0.004), indicating that they are quite homogeneous. Differentially methylated 100-kb windows were defined as those with false discovery rate (FDR)-adjusted p-value < 0.05 and absolute methylation change ≥ 5% relative to the mean for normal adrenal medulla.

**Figure 2.**
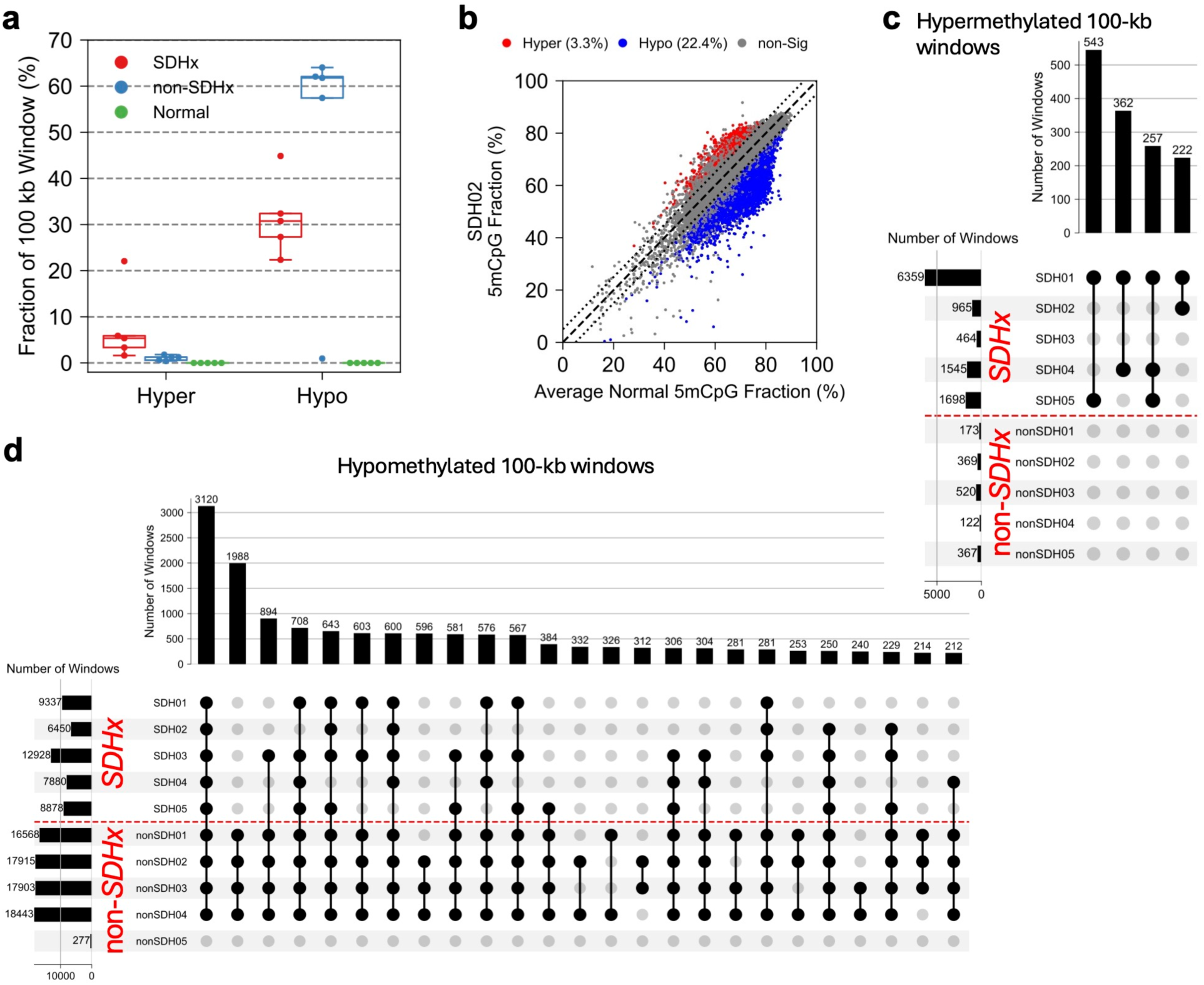
Global hypomethylation patterns overlap in *SDHx* and non-*SDHx* tumors. (**a**) Fractions of hyper- and hypo-methylated 100-kb windows in *SDHx* tumors and non-*SDHx* tumors relative to normal adrenal medulla cohort average. Boxplots for all autosomes (n = 5 per group). Dots indicate individual samples. Box elements as in Fig. 1a. (**b**) Scatterplot comparing methylation in 100-kb windows for SDH02 with the normal cohort average. Each dot represents a 100-kb window. Dashed line indicates equal methylation in SDH02 and the normal cohort average; dotted lines enclose windows differing in methylation by 5% or less. See Supplementary Fig. 3 for all samples. (**c**) UpSet plot for hypermethylated windows in tumors. The top bar plot shows the number of hypermethylated windows shared between samples. The side bar plot shows the number of hypermethylated windows in each tumor. Connected dots represent intersection groups. Only groups with at least two samples and 200 windows are shown. (**d**) UpSet plot for hypomethylated windows in tumors. Plot elements and filtering criteria as in (c).

This analysis again illustrates the bias towards hypomethylation in *SDHx* tumors (Fig. 2a). In the five *SDHx* tumors, 22.4% to 44.9% of 100-kb windows are hypomethylated, whereas only 1.6% to 22.1% of windows are hypermethylated. Four of the five non-*SDHx* tumors exhibit a large fraction of hypomethylated windows (57.4 to 64.0%) with almost no hypermethylated windows (≤ 2.0%), whereas nonSDH05 resembles normal adrenal medulla. Comparison of the methylation status of each 100-kb window in SDH02 with the same window in the normal adrenal medulla cohort using a scatter plot (Fig. 2b) confirms the preponderance of hypomethylated windows relative to hypermethylated windows. Scatter plots for all samples are provided in Supplementary Fig. 3.

To determine the extent to which the same 100-kb windows are affected in the different tumors, we selected differentially methylated windows that occur in at least two of the ten tumors and visualized groups containing at least 200 windows. There are relatively few hypermethylated windows in non-*SDHx* tumors, ranging from 122 to 520 windows, with little overlap among these tumors (Fig. 2c). There are many more hypermethylated windows in the *SDHx* tumors, ranging from 464 to 6,359, with some limited overlap between pairs of *SDHx* tumors (Fig. 2c). In contrast, hypomethylated windows show extensive overlap (Fig. 2d). Many of the hypomethylated windows present in the four globally hypomethylated non-*SDHx* tumors are not hypomethylated in *SDHx* tumors. However, nearly all recurrent hypomethylated windows in *SDHx* tumors are also hypomethylated in the four non-*SDHx* tumors that are globally hypomethylated (nonSDH01, 02, 03 and 04).

### Hypermethylated and hypomethylated CpG islands in *SDHx* tumors

Our observation that *SDHx* tumor DNA is hypomethylated apparently contradicts previous studies showing that *SDHx* mutations are associated with hypermethylated DNA ^7, 28^. However, these studies primarily interrogated CpG islands, rather than measuring DNA methylation genome-wide, as we have done. There are ∼27,000 CpG islands in the human genome (excluding the X and Y chromosomes). We compared the methylation of each CpG island in each normal sample with the average methylation of that island in all five normal samples using scatter plots (Fig. 3a). These plots are quite similar, except for Normal05, which exhibits a wider data spread. Most CpG islands segregate into two distinct subpopulations: hypo- and hyper-methylated, with relatively few islands methylated at intermediate levels (Fig. 3a; Fig. 4a, "all" islands).

**Figure 3.**
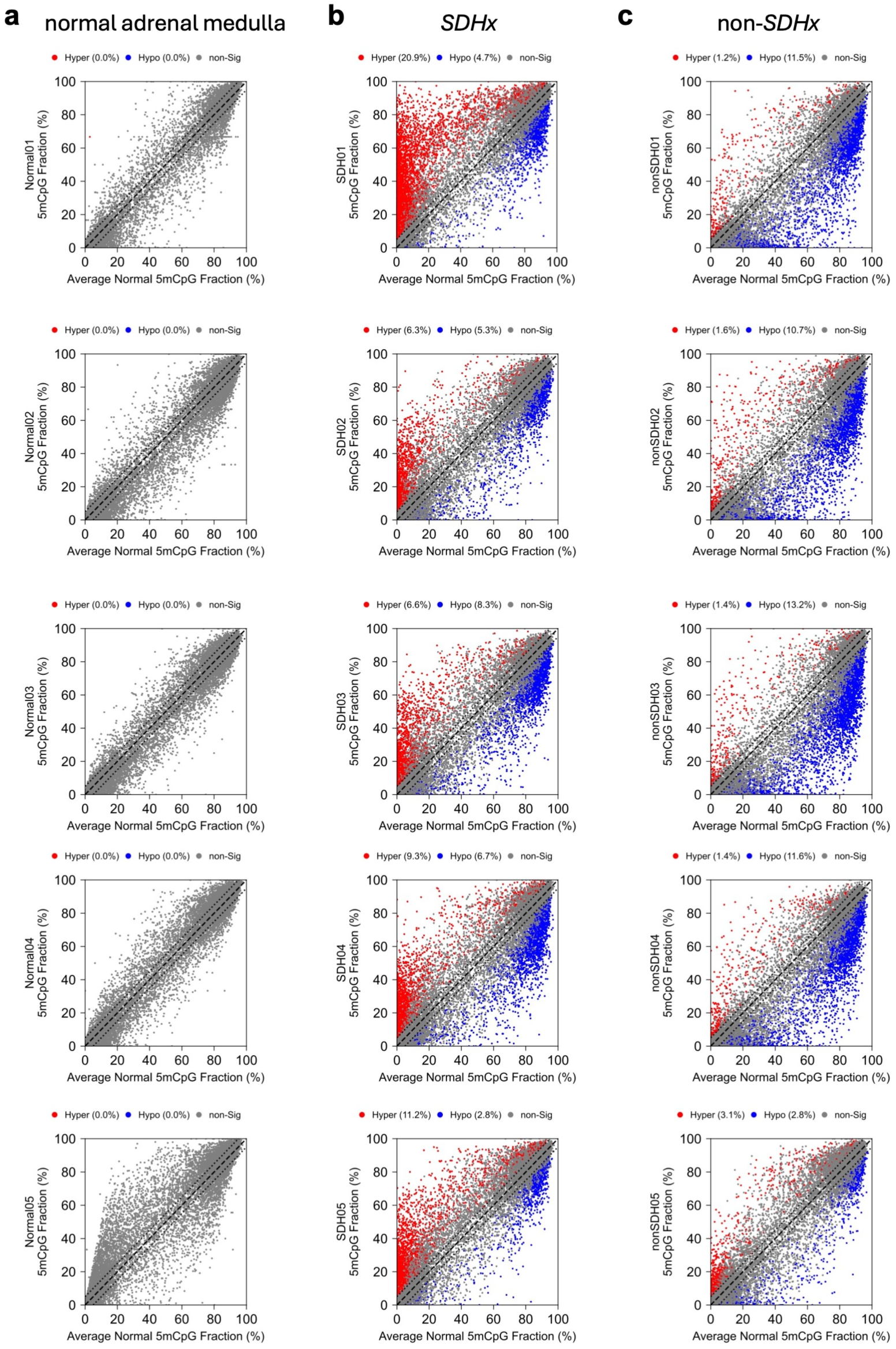
Hypermethylated and hypomethylated CpG islands in PPGL tumors. Scatter plots comparing methylation at all CpG islands in each sample with the normal cohort average (n = 5). Dashed line indicates equal methylation in the sample and the normal cohort average; dotted lines enclose islands differing in methylation by 5% or less. (**a**) Normal adrenal medulla. (**b**) *SDHx* tumors. (**c**) Non-*SDHx* tumors.

**Figure 4.**
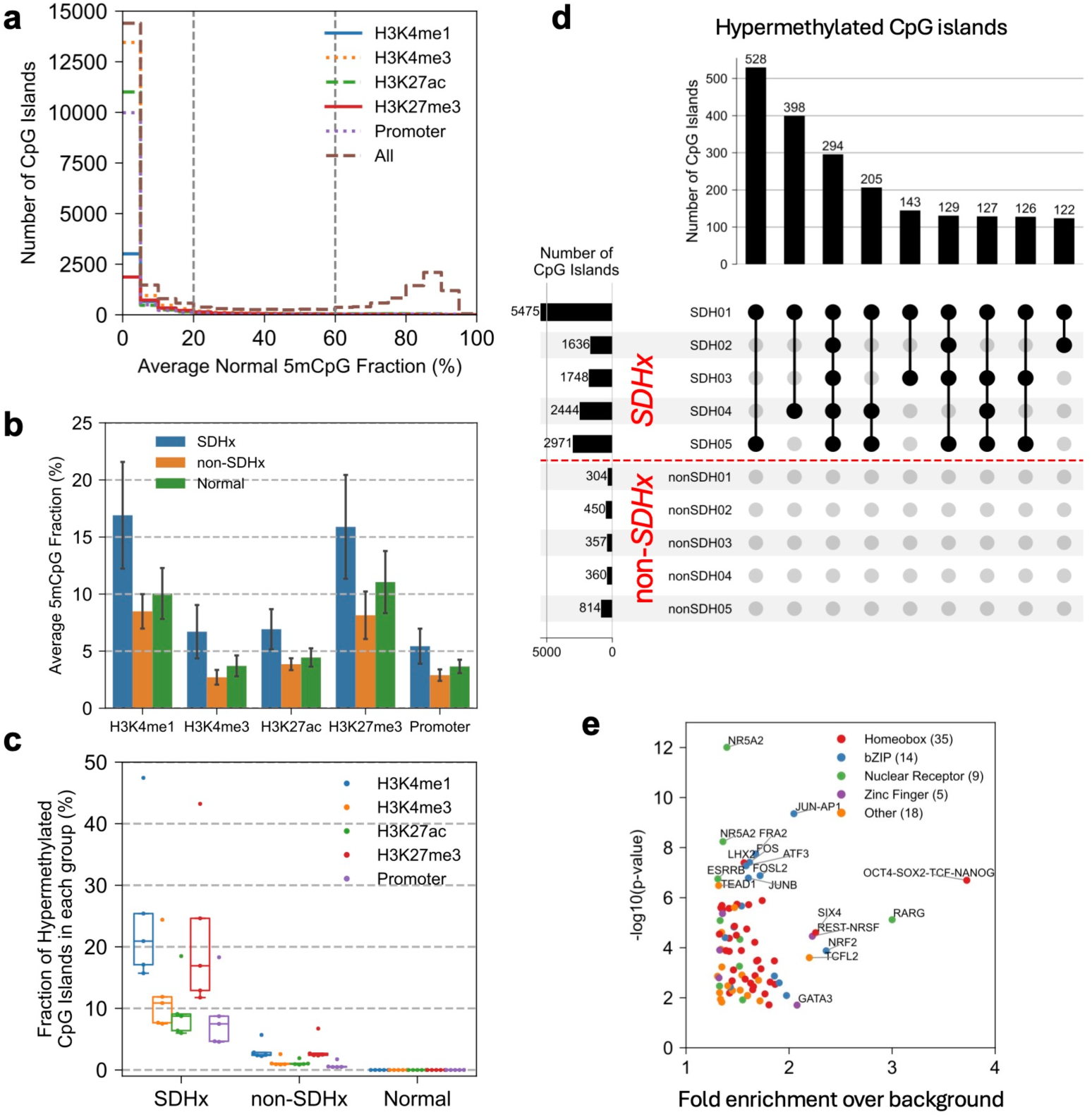
Histone-marked CpG islands tend to be hypermethylated in *SDHx* tumors. **(a)** Distribution of average DNA methylation levels for CpG islands in the normal cohort (n = 5). Number of CpG islands in each 5% methylation bin. CpG islands classified by overlap with histone modification peaks (ENCODE). Promoters: CpG islands overlapping the -1000 to +500 bp region relative to the TSS. (**b**) Average CpG island methylation by chromatin state. Bars represent the mean methylation of CpG islands in each chromatin state; error bars represent standard deviation (n = 5 per group). (**c**) Chromatin state of hypermethylated CpG islands. Box elements as in Fig. 1a. (**d**) Overlap of hypermethylated CpG islands in tumor samples. UpSet plot showing only intersection groups including at least two samples and 100 CpG islands. Plot elements as in Fig. 2c. (**e**) Enriched transcription factor motifs in hypermethylated CpG islands in *SDHx* tumors. Motif enrichment analysis using HOMER on CpG islands hypermethylated in at least three *SDHx* samples, with unchanged CpG islands as background control. Dots represent motifs with fold change > 1.3 and adjusted p-value < 0.05. Significantly enriched motifs (fold change > 2 or -log₁₀(p-value) > 6) are labeled.

We compared CpG island methylation in the tumors with normal adrenal medulla (Fig. 3). Although most CpG islands are unaffected in both types of tumor (similar % methylation in the tumor and in normal adrenal medulla; grey dots in Fig. 3), four of five *SDHx* tumors have more hypermethylated islands (6.3% to 20.9%) than hypomethylated islands (2.8% to 8.3%), relative to normal adrenal medulla (Fig. 3b). Non-*SDHx* tumors are more varied and differ from *SDHx* tumors: four of the five non-*SDHx* tumors show a bias toward hypomethylation of CpG islands; nonSDH05 shows small changes in both directions (Fig. 3c). NonSDH01, nonSDH02, nonSDH03 and nonSDH04 have many more hypomethylated islands (11.5%, 10.7%, 13.2% and 11.6%, respectively) than hypermethylated islands (1.2%, 1.6%, 1.4% and 1.4%, respectively). However, nonSDH05 has slightly more hypermethylated islands (3.1%) than hypomethylated islands (2.8%), although there are fewer hypermethylated islands in these tumors than in the *SDHx* tumors.

In summary, some, but not all, hypomethylated CpG islands in normal adrenal medulla are hypermethylated in *SDHx* tumors, whereas some hypermethylated islands in normal adrenal medulla are hypomethylated in *SDHx* tumors.

### Some but not all histone-marked CpG islands are hypermethylated in *SDHx* tumors

The chromatin state of gene regulatory elements such as CpG islands is an indicator of element activity in a particular tissue. We classified CpG islands based on histone modification data from the ENCODE adrenal gland dataset ^35^. In normal adrenal medulla, CpG islands bearing active chromatin marks (H3K4me1, H3K4me3 and H3K27ac) are predominantly methylated at low levels (< 20% methylation; 15,147 of 17,249 islands) (Fig. 4a). The large majority of these CpG islands overlap with H3K4me3 peaks (88%) and, to a lesser extent, with H3K27ac peaks (69%) and H3K4me1 peaks (25%). CpG islands bearing H3K27me3 (18%), a mark for Polycomb-repressed developmental promoters, are also hypomethylated in normal adrenal medulla, but most of these islands (75%) also overlap with an active histone mark, indicating that they are bivalent. H3K9me3 peaks, which are mostly associated with constitutive heterochromatin, show minimal overlap with CpG islands (0.2%). In contrast, CpG islands lacking any of these histone marks (n = 9,009) are hypermethylated (> 60% methylated; n = 6,432). Most of these unmarked CpG islands do not overlap with coding gene promoters (defined as -1 kb to +500 bp relative to the major transcription start site (TSS)) (Fig. 4a). In conclusion, the large majority of hypomethylated CpG islands in normal adrenal medulla are associated with active marks (H3K4me1, H3K4me3 and H3K27ac), whereas most hypermethylated CpG islands are not marked by any of these histone modifications.

We examined the relationship between CpG island methylation in the tumors and histone marks, noting that histone marks in normal adrenal medulla and the tumors may be different. On average, CpG islands marked with H3K4me1, H3K4me3, H3K27ac or H3K27me3 in normal adrenal medulla are hypermethylated in *SDHx* tumors, relative to normal adrenal medulla and non-*SDHx* tumors (Fig. 4b). Although histone-marked islands are hypermethylated on average, the absolute methylation is still low (∼10% methylated on average; the genomic average methylation is ∼64%). In *SDHx* tumors, a larger fraction of hypermethylated CpG islands is associated with each histone mark than in non-*SDHx* tumors (Fig. 4c). This is true for both active and inactive histone marks, as well as for promoters with overlapping CpG islands. CpG islands marked by H3K4me1 or H3K27me3 are more likely to be hypermethylated in *SDHx* tumors, with ∼20% of islands hypermethylated compared to ∼10% for H3K4me3-, H3K27ac- and promoter-associated islands.

Scatter plot analysis of the individual samples shows that a minority of CpG islands marked with H3K4me1 and/or H3K4me3 are hypermethylated in *SDHx* tumors, ranging from 8% to 25% of these islands (Supplementary Fig. 4). This fraction is much lower in non-*SDHx* tumors (1% to 3%; Supplementary Fig. 4). In contrast, very few H3K4me1/3-marked islands are hypomethylated in *SDHx* tumors (1% or less) and non-*SDHx* tumors (< 4%) (Supplementary Fig. 4). Scatter plot analysis of unmarked CpG islands shows that relatively few of these islands are hypermethylated in *SDHx* tumors (up to 10%) and non-*SDHx* tumors (< 3%) (Supplementary Fig. 5). The fractions of unmarked hypomethylated islands are similar: *SDHx* tumors (7% to 24%); non-*SDHx* tumors (6% to 34%) (Supplementary Fig. 5).

In summary, the methylation status of most CpG islands is unchanged in *SDHx* tumors relative to normal adrenal medulla. However, a varying fraction of CpG islands is hypermethylated in these tumors, consistent with succinate inhibition of demethylation of some islands, but not all.

### Enriched transcription factor motifs in *SDHx*-associated hypermethylated CpG islands

Hypermethylated CpG islands are often shared among *SDHx* tumors, whereas non-*SDHx* tumors have relatively few hypermethylated islands with no overlap with other tumors (Fig. 4d). To identify potential regulatory mechanisms, we used HOMER to perform motif enrichment analysis on CpG islands that are hypermethylated in at least three *SDHx* tumors (n = 2,030; maximum length = 5 kb). CpG islands showing no significant change in methylation were used as background (n = 16,462; maximum length = 5 kb). We identified 81 enriched motifs (q < 0.05, fold enrichment > 1.3; Fig. 4e), including homeobox (n = 35), bZIP (n = 14), nuclear receptor (n = 9) and zinc finger (n = 5) transcription factor families (other families n = 18). The most significantly enriched motifs (-log₁₀(p-value) > 6) include AP-1 family members (JUN-AP1, FRA2, FOS, ATF3 and JUNB), which regulate stress response and cellular adaptation, and nuclear receptors (NR5A2 and ESRRB), which are involved in metabolic regulation. Motifs with the highest fold enrichment (> 2) include the pluripotency factors (OCT4-SOX2-TCF-NANOG; 3.7-fold), retinoic acid receptor RARG (3.0-fold) and the neural repressor REST-NRSF (2.2-fold). Although these enrichments are small, they may hint at altered gene regulatory mechanisms in *SDHx* tumors.

### CpG island promoters are not generally more methylated in *SDHx* tumors

CpG islands often overlap regulatory elements, such as enhancers and promoters. We divided coding gene promoters into two groups, depending on whether they overlap a CpG island. Promoters overlapping CpG islands (n = 12,473) are almost completely demethylated around the TSS in normal adrenal medulla (Fig. 5a, upper panels). In *SDHx* tumors, these promoters are only marginally more methylated than in normal adrenal medulla, with SDH01 showing the largest effect. In non-*SDHx* tumors, these promoters are methylated at similar levels to normal adrenal medulla near the TSS. However, relative to the normal adrenal medulla average, nonSDH01, nonSDH02, nonSDH03 and nonSDH04 are hypomethylated in the flanking regions (these are the globally hypomethylated tumors).

**Figure 5.**
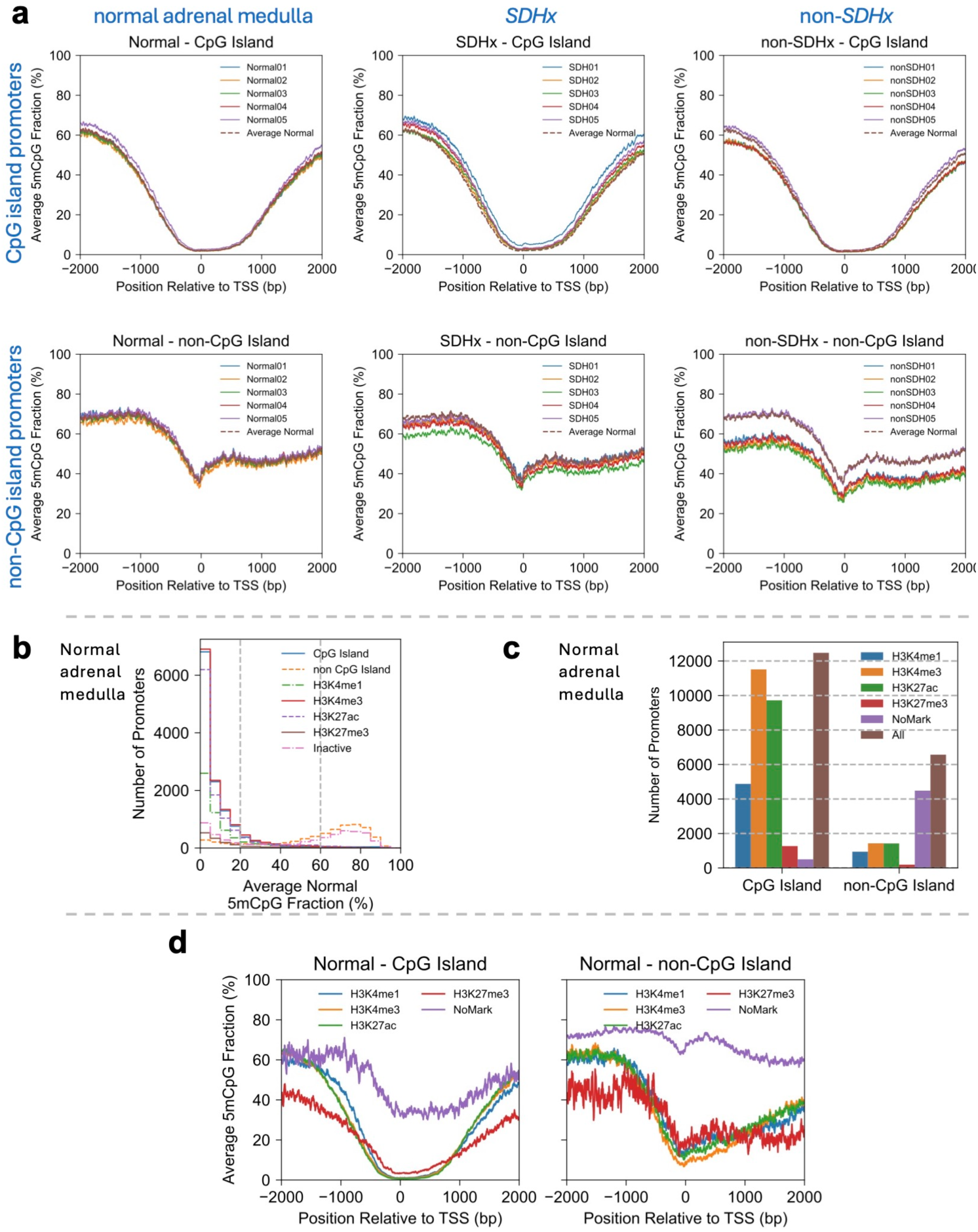
CpG island promoters are not generally hypermethylated in *SDHx* tumors. (**a**) CpG island and non-island promoters: Average DNA methylation profiles relative to the TSS for normal adrenal medulla, *SDHx* tumors and non-*SDHx* tumors. (**b**) Distribution of average promoter methylation levels in the normal adrenal medulla cohort (n = 5). The y-axis shows the number of promoters in each 5% methylation bin. (**c**) Number of CpG island and non-island promoters with specific histone marks. “All”: all CpG island promoters or non-CpG island promoters. "No Mark": promoters with none of the listed histone marks. (**d**) Average DNA methylation profiles relative to the TSS for CpG island and non-island promoters marked with H3K4me1, H3K4me3, H3K27ac or H3K27me3 in normal adrenal medulla. Promoters were classified by overlap with CpG islands and histone modification peaks from ENCODE.

Promoters that do not overlap with CpG islands (n = 6,583) are much less hypomethylated around the TSS (Fig. 5a, lower panels). The methylation levels in normal adrenal medulla are very similar in all five samples. The *SDHx* tumors are similar to the normal adrenal medulla average, except for SDH03, which is hypomethylated relative to the others. The non-*SDHx* tumors are more varied, such that nonSDH01, nonSDH02, nonSDH03 and nonSDH04 are again hypomethylated relative to the others and to normal adrenal medulla.

We correlated promoter DNA methylation with histone marks in normal adrenal medulla. Almost all CpG island promoters are hypomethylated and marked with H3K4me3, H3K27ac and, to a lesser extent, with H3K4me1 (Fig. 5b). In contrast, most, but not all, non-CpG island promoters are hypermethylated and lack histone marks (Fig. 5b). More generally, CpG island promoters are enriched in active marks relative to non-CpG island promoters (Fig. 5c). CpG island promoters with active histone marks (H3K4me1, H3K4me3 and H3K27ac) are completely demethylated around the TSS (Fig. 5d, left panel). CpG island promoters with the inactive H3K27me3 mark (mostly bivalent) are also demethylated around the TSS but differ from those with active marks in that their flanking DNA is hypomethylated. CpG island promoters lacking any of these marks are only mildly hypomethylated around the TSS relative to the genomic average (∼68% methylation). Non-island promoters show similar trends, but to a lesser extent (Fig. 5d, right panel): active marks correlate with hypomethylation around the TSS, the bivalent mark (H3K27me3) correlates with hypomethylated flanking DNA, and unmarked promoters are only mildly hypomethylated.

In summary, in normal adrenal medulla, CpG island promoters tend to be extremely hypomethylated around the TSS and carry active histone marks. In *SDHx* tumors, the average CpG island promoter methylation is only slightly higher than in normal adrenal medulla, because the fraction of hypermethylated CpG island promoters is relatively small.

### Active promoters are hypomethylated relative to inactive promoters

To assess the potential functional relevance of promoter methylation differences, we compared our data with average gene expression levels from the TCGA pheochromocytoma and paraganglioma (PCPG) cohort (n = 179 primary tumor samples ^5^). Although matched expression data are unavailable for our samples, TCGA provides representative transcriptional profiles for this tumor type. We classified inactive genes (TPM < 1) and stratified active genes into expression quintiles (Ǫ1 to Ǫ5, in ascending order of expression level), and examined their promoter methylation patterns (Fig. 6a). Inactive promoters are much more methylated (averaging ∼ 45% methylated) than active promoters (< ∼20% methylation) in normal adrenal medulla and both types of tumor. In addition, higher transcriptional activity correlates with decreasing promoter methylation (from ∼20% to ∼10% average methylation) in normal adrenal medulla and both tumors (Fig. 6a).

**Figure 6.**
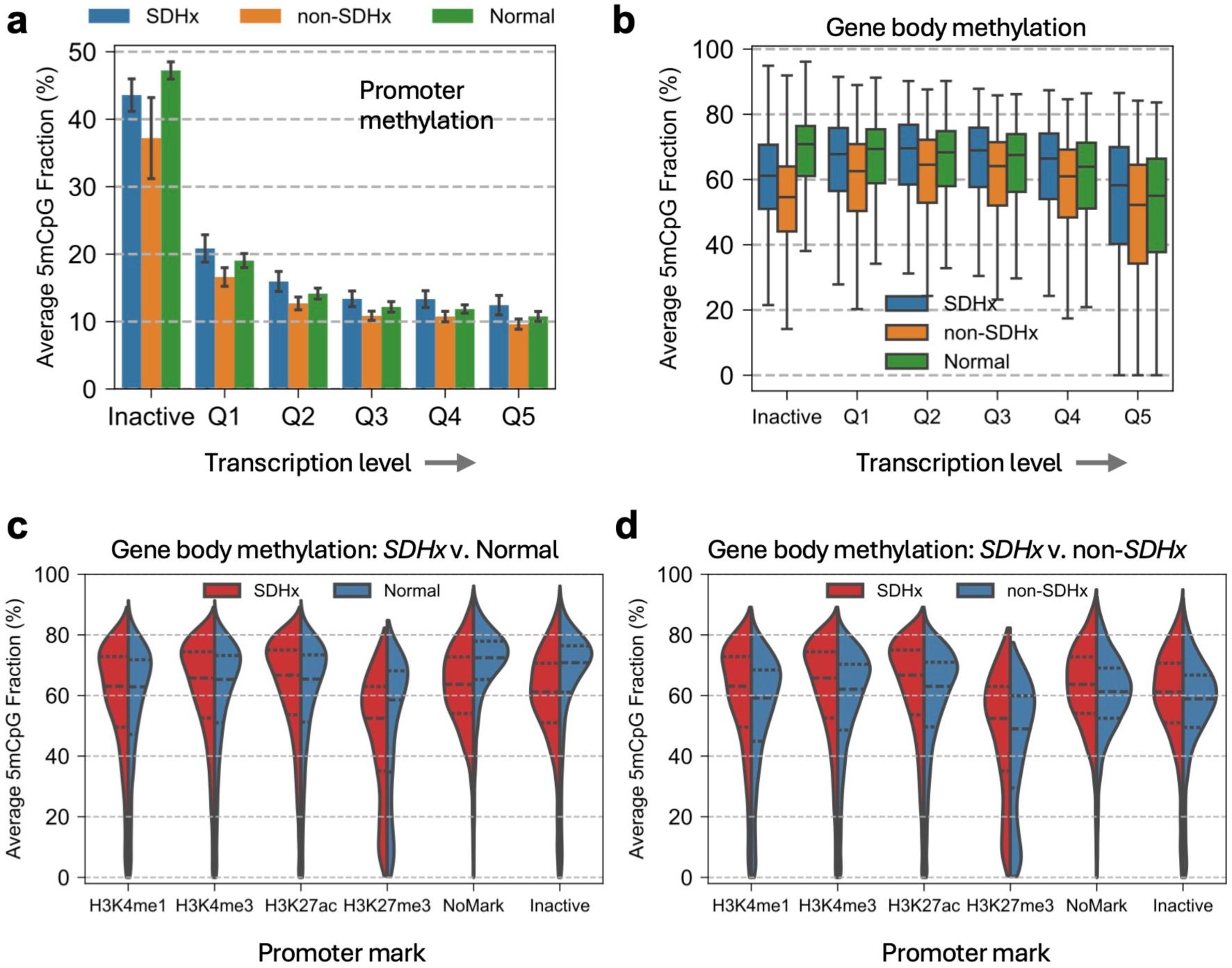
Inactive gene bodies tend to be hypomethylated in both PPGL tumor types. (**a**) DNA methylation status of active and inactive promoters. Bar plot of average promoter methylation by expression level and sample group. Protein-coding genes were partitioned into inactive genes and increasingly active genes denoted by quintiles Ǫ1-Ǫ5. Average expression data were obtained from the TCGA-PCPG cohort (n = 179). Error bars represent standard deviation (n = 5 per group). (**b**) DNA methylation status of active and inactive gene bodies. Average gene body methylation sorted by expression level and sample group. Box elements as in Fig. 1a. (**c**) Gene body methylation sorted by promoter histone mark in normal adrenal medulla and *SDHx* tumors (violin plots). Black dashed lines represent the 25th, 50th and 75th percentiles of data in each group. (**d**) As in (c), but comparing gene body methylation in *SDHx* and non-*SDHx* tumors.

In general, active and inactive gene bodies average more methylated than their promoters, particularly for active genes (Fig. 6a,b). Active gene bodies show similar but weaker trends relative to their promoters: decreasing methylation with increasing activity. Inactive gene bodies are hypomethylated in both tumors relative to normal adrenal medulla (Fig. 6b). We classified gene bodies according to their promoter histone marks (Fig. 6c). Gene bodies with active histone marks at their promoters (H3K4me1, H3K4me3 and H3K27ac) exhibit similar methylation patterns in *SDHx* tumors and normal adrenal medulla, although the distributions are shifted slightly toward hypermethylation in *SDHx* tumors (Fig. 6c). However, gene bodies with promoters bearing the inactive H3K27me3 mark are hypomethylated in *SDHx* tumors relative to normal adrenal medulla. They include a relatively large fraction of lowly methylated gene bodies, which might have bivalent histone marks. Genes with promoters lacking a histone mark are hypomethylated in *SDHx* tumors, corresponding mostly to inactive genes. Gene bodies are generally hypomethylated in non-*SDHx* tumors relative to *SDHx* tumors and normal adrenal medulla (Fig. 6c,d).

### Centromeric elements are exceptionally hypomethylated in both PPGL tumor types

We examined methylation at repetitive sequences: LINE elements (n = 368,904), SINE elements (n = 921,534) and LTR retrotransposons (n = 255,848), which together comprise about 44% of the genome. These repetitive elements are highly methylated in normal adrenal medulla, with median methylation levels of 76.3% for LINEs, 84.2% for SINEs and 76.0% for LTRs (Fig. 7a). In contrast, both *SDHx* and non-*SDHx* tumors show widespread hypomethylation across all repeat classes, with median decreases ranging from 2.1 to 11.4% (Fig. 7b). The magnitude of hypomethylation is comparable across repeat types in both *SDHx* and non-*SDHx* tumors. All three element types average less hypomethylated in *SDHx* tumors than in non-*SDHx* tumors (Fig. 7b).

**Figure 7.**
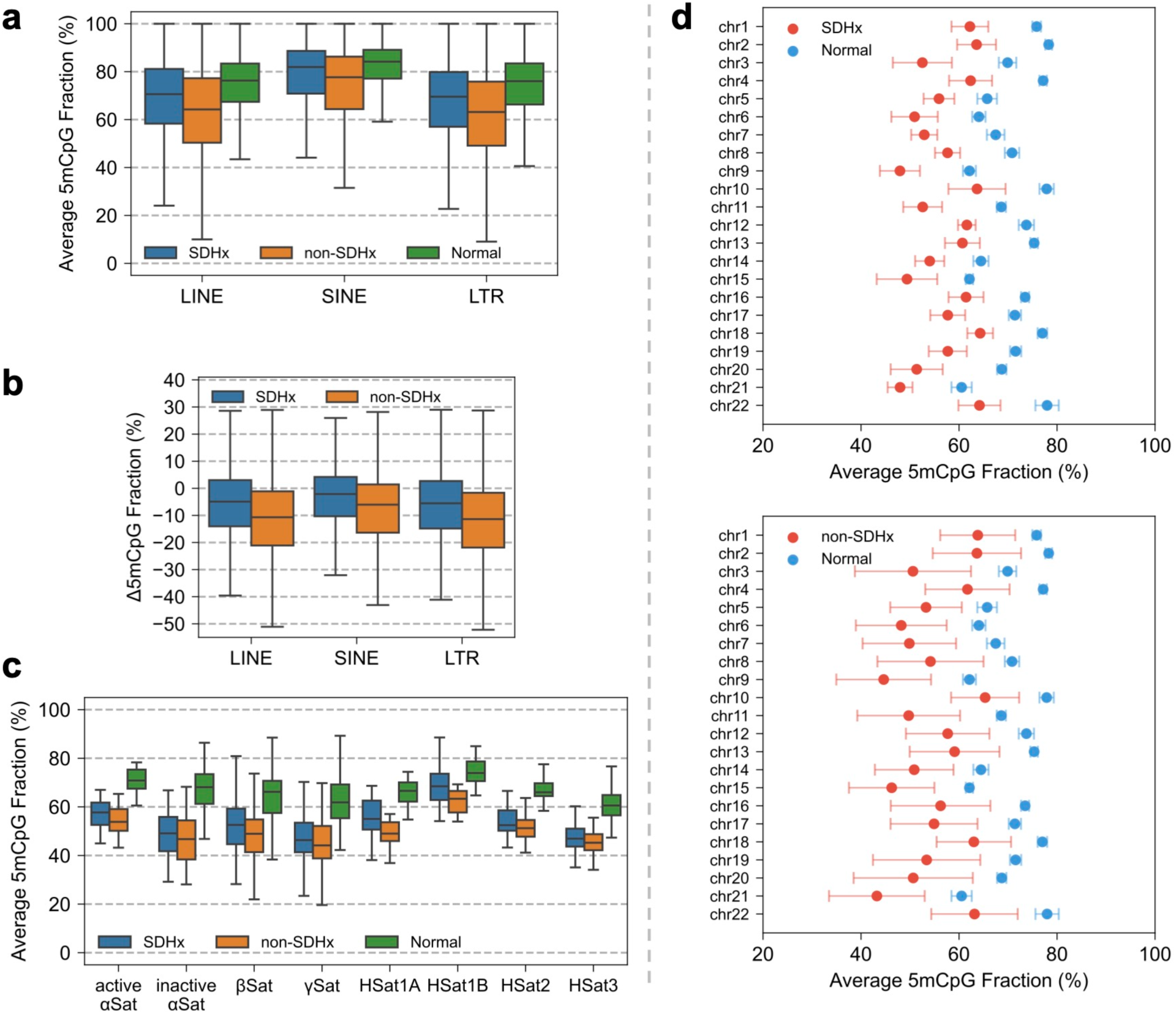
Centromeric elements are exceptionally hypomethylated in PPGL tumors. **(a)** Average methylation of LINE, SINE and LTR repetitive elements in normal adrenal medulla and the tumors. Box elements as in Fig. 1a. **(b)** Differential methylation of repetitive elements. Distribution of methylation differences (tumor minus normal) for individual repetitive elements. **(c)** Average methylation of centromeric elements in normal adrenal medulla and the tumors. Boxplot of average methylation by centromeric element class and sample group. Box elements as in Fig. 1a. **(d)** Point-range plots comparing methylation levels in active α-satellites between normal adrenal medulla and *SDHx* tumors (upper panel) or non-*SDHx* tumors (lower panel). Points represent the mean methylation of active α-satellites in each autosome (5 samples per group; error bars: standard deviation).

Centromeric elements display a much more pronounced hypomethylation than the other repetitive sequences (Fig. 7c). Active α-satellite repeats, which maintain centromere identity and chromosome segregation, have a median methylation of 70.8% in normal adrenal medulla, but only 57.7% and 53.8% in *SDHx* and non-*SDHx* tumors, respectively. This exceptional hypomethylation is observed across all autosomes in both types of tumor (Fig. 7d), with a mean difference in the methylation level of each chromosome of -13.7 ± 1.8% for *SDHx* tumors and -15.9 ± 2.1% for non-*SDHx* tumors (SD; n = 22 autosomes). These data show that centromeres in PPGL tumors tend to be extremely hypomethylated, although this effect is unlikely to be mediated by succinate since both *SDHx* and non-*SDHx* tumors are affected.

## Discussion

In summary, we have shown that *SDHx* tumors are globally hypomethylated and therefore similar to most tumors in this regard. In particular, non-promoter CpG islands, inactive gene bodies, repetitive sequences and centromeres are all hypomethylated in *SDHx* tumors. *SDHx* tumors differ from non-*SDHx* tumors in that a much larger fraction of CpG islands is hypermethylated (6% - 21% vs. 1% - 3%). The difference between our study and those reported previously is that we used long-read sequencing for direct detection of DNA methylation genome-wide, whereas they used microarray approaches focused on CpG islands ^28^. Ideally, we would have examined many more than five *SDHx* tumors; however, nanopore sequencing of the complete human genome is very costly. We also note that different *SDHx* mutations may alter SDH enzyme activities and succinate sensitivity to different extents (e.g., SDH01 shows stronger effects than the other *SDHx* tumors) ^36^. Nevertheless, our data are quite consistent. Clearly, the simplest model, that succinate inhibits TET DNA demethylases resulting in global hypermethylation, does not account for our data.

CpG islands are methylated by the de novo DNMT3A/B DNA methylases ^37^, which are inhibited by H3K4me1/3 ^38, 39, 40, 41, 42^. Both enzymes contain an ADD domain that functions as a sensor for H3K4 methylation. When H3K4 is unmethylated, the ADD domain binds the H3 tail, freeing the catalytic domain to methylate DNA. When H3K4 is methylated, the ADD domain folds back onto the catalytic domain, blocking it from binding DNA. As a result, regions marked by H3K4 methylation are shielded from de novo DNA methylation. We found that, in normal adrenal medulla, 61% of all autosomal CpG islands are marked by H3K4me1/3 and methylated at very low levels, consistent with their role as active gene regulatory elements. This low level of DNA methylation can be accounted for by H3K4me1/3 inhibition of DNMT3A/B, combined with TET DNA demethylase activity (Fig. 8a). In contrast, CpG islands lacking H3K4me1/3 are mostly highly methylated and inactive, presumably because DNMT3A/B are not inhibited and therefore more active than the TET DNA demethylases (Fig. 8a).

**Figure 8.**
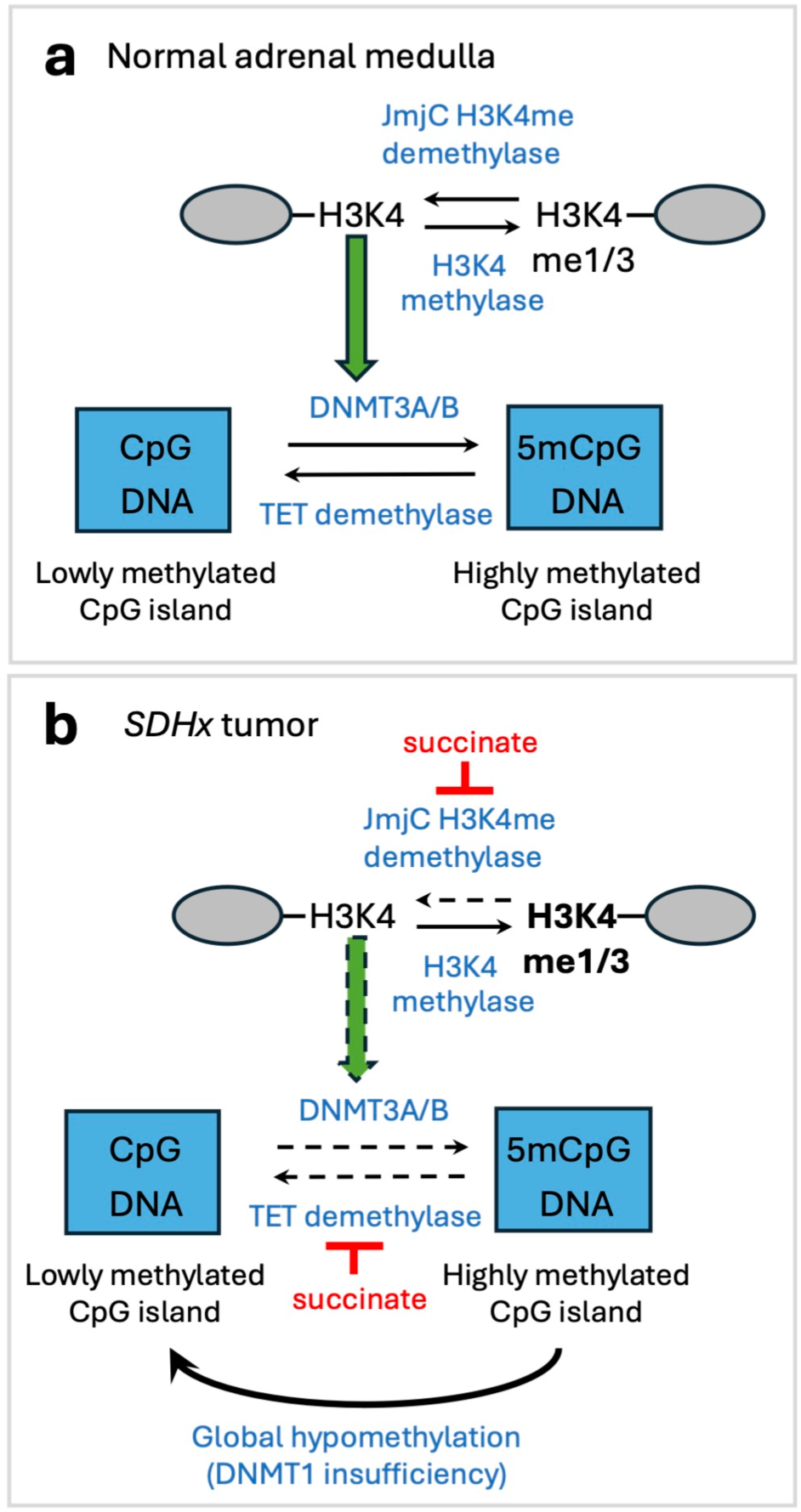
Model for succinate effects on CpG island methylation in *SDHx* tumors. (**a**) In normal tissue, such as adrenal medulla, the level of CpG island methylation is determined by the balance between DNMT3A/B and TET activities. Since DNMT3A/B is activated by unmethylated H3K4, this balance will be determined by the extent of H3K4 methylation. If H3K4 is unmethylated, the CpG island will be highly methylated. If H3K4 is methylated, DNMT3A/B activity is low, TET activity is unaffected, and the island will be lowly methylated. Island targeting is achieved through recruitment of H3K4 methylase activity by sequence-specific transcription factors. Grey oval: nucleosome. (**b**) In *SDHx* tumors, succinate inhibits JmjC H3K4me demethylases, resulting in increased H3K4me, and reduced DNMT3A/B activity, tending to reduce CpG island methylation. However, direct inhibition of TET demethylases by succinate results in the opposite effect, tending to increase DNA methylation. The extent of island methylation will reflect the balance between these opposing succinate effects, overlaid on the global DNA hypomethylation presumed due to DNMT1 insufficiency. Dashed arrows indicate reduced activity.

Four of the five non-*SDHx* tumors are globally hypomethylated, similar to other tumors. However, we find that relatively few CpG islands (< 4% of islands) marked by H3K4me1/3 are hypomethylated in non-*SDHx* tumors, and that a similarly small fraction (< 3% of islands) is hypermethylated, indicating that H3K4me1/3-marked islands are not strongly affected by the global hypomethylation mechanism acting in tumors. On the other hand, a larger fraction of unmarked CpG islands is hypomethylated in non-*SDHx* tumors (25% to 34% of islands, except for nonSDH05 with 6%), whereas only a small fraction is hypermethylated (< 3% of islands). A potential explanation is that CpG islands marked by H3K4me1/3 are already hypomethylated (4% to 7% methylation) in normal adrenal medulla, whereas those lacking this histone mark are mostly hypermethylated (62% to 66% methylation) in normal adrenal medulla, resulting in more obvious hypomethylation of unmarked islands in these tumors.

In *SDHx* tumors, succinate inhibition of TET DNA demethylases would tend to counter the global hypomethylation typical of most tumors, predicting a general hypermethylation relative to non-*SDHx* tumors. However, we observed global hypomethylation in *SDHx* tumors similar to that observed in other tumors, suggesting that the succinate effect on TET DNA demethylases is weaker than the mechanisms inhibiting maintenance methylation. We have shown that hypermethylation in *SDHx* tumors is restricted to a subset of CpG islands, suggesting that the TET DNA demethylases might be targeted to these islands. However, there is little evidence for targeting of TET DNA demethylases (although the sequence context of the CpG affects the demethylation rate ^43^), or for their inclusion in targeted protein complexes similar to those in which histone modifying enzymes are usually found. If this is true, another explanation for targeted succinate-associated hypermethylation at some CpG islands is needed.

Unlike the TET DNA demethylases and the DNMT1 DNA methylase, the de novo DNMT3A/B DNA methylases are targeted to CpG islands ^44, 45^. Consequently, we propose that indirect succinate inhibition of targeted DNMT3A/B DNA methylases is the key (Fig. 8b). The JmjC H3K4me histone demethylases are also inhibited by succinate since they use the same enzymatic mechanism as the TET DNA demethylases ^46^. Consequently, an increase in succinate concentration would inhibit both the TET DNA demethylases (predicting hypermethylation) and the H3K4me1/3 demethylases (inhibiting DNMT3A/B methylases, predicting hypomethylation). Thus, the actual DNA methylation level may depend on how succinate shifts the balance between these two activities at each CpG island.

At CpG islands lacking H3K4me1/3, DNMT3A/B enzymes are active, whereas TET demethylases are inhibited by succinate, predicting hypermethylation (Fig. 8b). However, more unmarked islands are hypomethylated than hypermethylated in each *SDHx* tumor, which is also true for the non-*SDHx* tumors. To explain this effect, we propose that, at CpG islands lacking H3K4me1/3, the hypomethylation effect due to impaired maintenance methylation by DNMT1 dominates over the hypermethylation effect of TET DNA demethylase inhibition by succinate, shifting the absolute methylation level lower (Fig. 8b). In contrast, at CpG islands marked by H3K4me1 and/or H3K4me3, the inferred effect of succinate is clear: 8% to 25% of these islands are hypermethylated, whereas very few (< 1%) are hypomethylated. The implication is that, at these islands, the inhibitory effect of succinate on TET DNA demethylases is greater than its inhibitory effect on DNMT3A/B through the JmjC histone demethylases, shifting the balance towards increased DNA methylation, and overcoming global hypomethylation (Fig. 8b).

Finally, we note that our observations may help to explain the lack of *SDHx* tumor response to a DNA methylase inhibitor in a clinical trial ^47^. Our results suggest that a DNA methylase inhibitor might exacerbate global hypomethylation as well as reducing CpG island methylation. In addition, the magnitude of centromeric hypomethylation - approximately two-fold greater than at other repetitive elements - suggests vulnerability to chromosome instability in both *SDHx* and non-*SDHx* PPGL tumors ^48^, since global DNA hypomethylation is associated with chromosomal instability in cancer cells ^49, 50^.

## Methods

### DNA extraction, nanopore library preparation and sequencing

Frozen tissue samples were carefully sliced on dry ice. Genomic DNA was extracted from 15–25 mg of tissue using the PureLink Genomic DNA Mini Kit (Invitrogen K182002) or from 0.5–1.0 ml whole blood using the Puregene Blood Kit (Ǫiagen 158389), following the manufacturers’ protocols. DNA concentrations and purity were quantified by measuring A260 and the A260/A280 ratio. Sequencing libraries were prepared as described ^51, 52^. Briefly, 1.2–2.0 µg of genomic DNA was used for nanopore library preparation. DNA repair and end preparation were performed using the NEBNext Companion Module for ONT (New England Biolabs E7180L), followed by adapter ligation, flow cell priming and library loading using kit V14 (Oxford Nanopore Technologies (ONT) SǪK-LSK114) according to the manufacturer’s instructions. Each sample was sequenced on a separate flow cell (ONT R10 version FLO-MIN114) using either a MinION Mk1C or a GridION instrument (ONT). To maximize sequencing yield, a fraction of each library was first loaded onto the flow cell and sequenced for 48-72 h. Each flow cell was then washed at least once using the ONT Flow Cell Wash Kit (EXP-WSH004), including DNase I treatment, and reloaded with the remaining library for a further 48-72 h of sequencing.

### Nanopore sequencing

Oxford Nanopore reads were base-called using Dorado v1.0.0 with the dna_r10.4.1_e8.2_400bps_sup v5.2.0 base-calling model and 5mCG_5hmCG v1 modification model (--min-qscore 8). Reads were aligned to the T2T-CHM13v2.0 assembly (with pseudo-autosomal regions (PARs) on Y masked to N and with rCRS (AC:NC_012920.1) sequence for mitochondrial DNA) ^32, 53^. 5mCpG and 5hmCpG sites were called using modkit v0.5.0 (--cpg -- combine-strands); the average 5mCpG methylation in each region was also computed with modkit stats with minimum coverage of 3 reads (-m 3).

### Detection of differentially methylated regions (DMRs) using a beta-binomial framework

We developed a beta-binomial framework to account for biological variability in the normal cohort and identify DMRs. In this framework, the number of mCpG sites (given total valid CpG sites from modkit) in normal tissue follows a binomial distribution, while the methylation rate follows a beta distribution with shape parameters *α* and *β* across the normal cohort. For each region in the normal cohort, we filtered for regions with at least three valid normal samples (filter criteria as in Nanopore sequencing) for analysis. Parameters (*α*, *β*) were estimated using maximum likelihood optimization implemented in custom Python code. The intraclass correlation coefficient (*ρ*) was calculated from the beta-binomial parameters as *ρ* = 1 / (1 + *α* + *β*), which quantifies the degree of overdispersion relative to a binomial model. We calculated two-tailed p-values and Benjamini-Hochberg adjusted p-values to control false discovery rates. Regions with adjusted p-values < 0.05 and absolute methylation changes ≥ 5% were classified as significant DMRs. Hyper- and hypo-methylated DMRs are defined by direction of methylation change. To ensure high-confidence predictions we retained only regions where at least three normal samples had enough coverage for at least 10 valid CpGs in the region.

### Computational analysis

For genome-wide analysis, we partitioned the 22 autosomes into non-overlapping 100 kb windows using custom python code. We used the JHU RefSeqv110 + Liftoff v5.2 annotation for protein-coding genes. Promoter regions were defined as -1 kb to +500 bp relative to the TSS. Gene bodies were defined as the region from the TSS to the Transcription End Site (TES). CpG islands were predicted using the UCSC cpg_lh command with default parameters ^54^. We used LINEs, SINEs, and LTR annotations from RepeatMasker v4.1.2p1.2022Apr14 and Cen/Sat v2.1 annotation for centromeric elements. We obtained histone modification peaks for hg38 assembly from the ENCODE adrenal gland dataset (ENCFF020HRP, ENCFF816NIJ, ENCFF817UME, ENCFF876NSG and ENCFF987SRC) ^35^. Peak locations in the T2T assembly were computed with UCSC LiftOver ^55^. We classified CpG islands and promoters as overlapping each histone mark if they had ≥ 30% overlap using bedtools (bedtools intersect -f 0.3) ^56^. We obtained RNA-seq data from the TCGA PCPG cohort ^5^ using TCGAbiolinks (R source code available on Figshare) ^57^. We selected only "Primary solid tumor" samples (n = 179) from the PCPG cohort; only protein-coding genes were analyzed. Active genes were defined as genes with TPM ≥ 1 in ≥ 50% of TCGA samples; all other genes were classified as inactive. Active genes were partitioned into Ǫ1-Ǫ5 quintiles by ascending expression level. All processed genomic annotations (CpG islands, 100 kb windows, histone mark classifications and gene expression classifications) are available at Figshare. For motif enrichment analysis, we identified shared hypermethylated CpG islands that were hypermethylated in at least three *SDHx* tumor samples (n = 2,030; maximum length = 5 kb). The background set consisted of shared non-significant CpG islands (n = 16,462; maximum length = 5 kb). We used HOMER v5.1 ^58^ to identify enriched motifs in hypermethylated CpG islands (-size given -mask -cpg) compared to non-significantly changed CpG islands, and filtered motifs for enrichment > 1.3 and adjusted p-values < 0.05.

## Data availability

The nanopore sequencing data will be available under controlled access at the NIH Database of Genotypes and Phenotypes (dbGaP), because the data may potentially include identifying information.

## Code availability

DOI for publication: 10.6084/m9.figshare.34754907

## Acknowledgements

This study used the high-performance computational capabilities of the Biowulf Linux cluster at the NIH.

## Author contributions

K.P., I.H. and H.A. obtained the patient samples; H.P., C.C. and P.E. performed the nanopore sequencing. Z.X. did the bioinformatic analysis. Z.X. and D.C. wrote the manuscript with input from all authors.

## Funding

This research was supported by the Intramural Research Program of the National Institutes of Health (NIH). The contributions of the NIH authors are considered Works of the United States Government. The findings and conclusions presented in this paper are those of the authors and do not necessarily reflect the views of the NIH or the U.S. Department of Health and Human Services.

## Competing interests

The authors declare no competing interests, except that Dr. Pacak is the Chair of the Medical Advisory Board, Pheo Para Alliance. He has also received honoraria from the American Association of Clinical Endocrinology.

## Additional information

Supplementary Figures 1 - 5 and Supplementary Table 1.

