## Supplemental Information for "Long-read DNA sequencing reveals global hypomethylation and focal hypermethylation in *SDHx*-deficient pheochromocytoma and paraganglioma"

#### Supplementary Figures 1-5.

#### Supplementary Table 1.

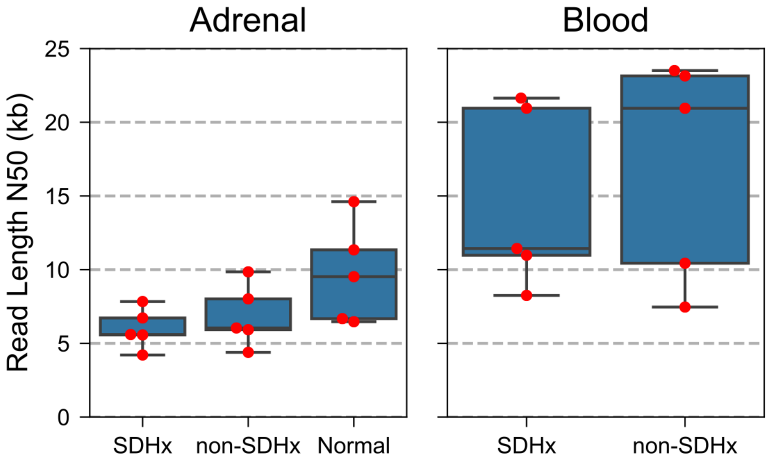


**Supplementary Fig. 1**. **Nanopore read N50 values**. Box boundaries show the 25th and 75th percentiles, the central line indicates the median, and whiskers extend to 1.5x the interquartile range or the most extreme data point within that range.

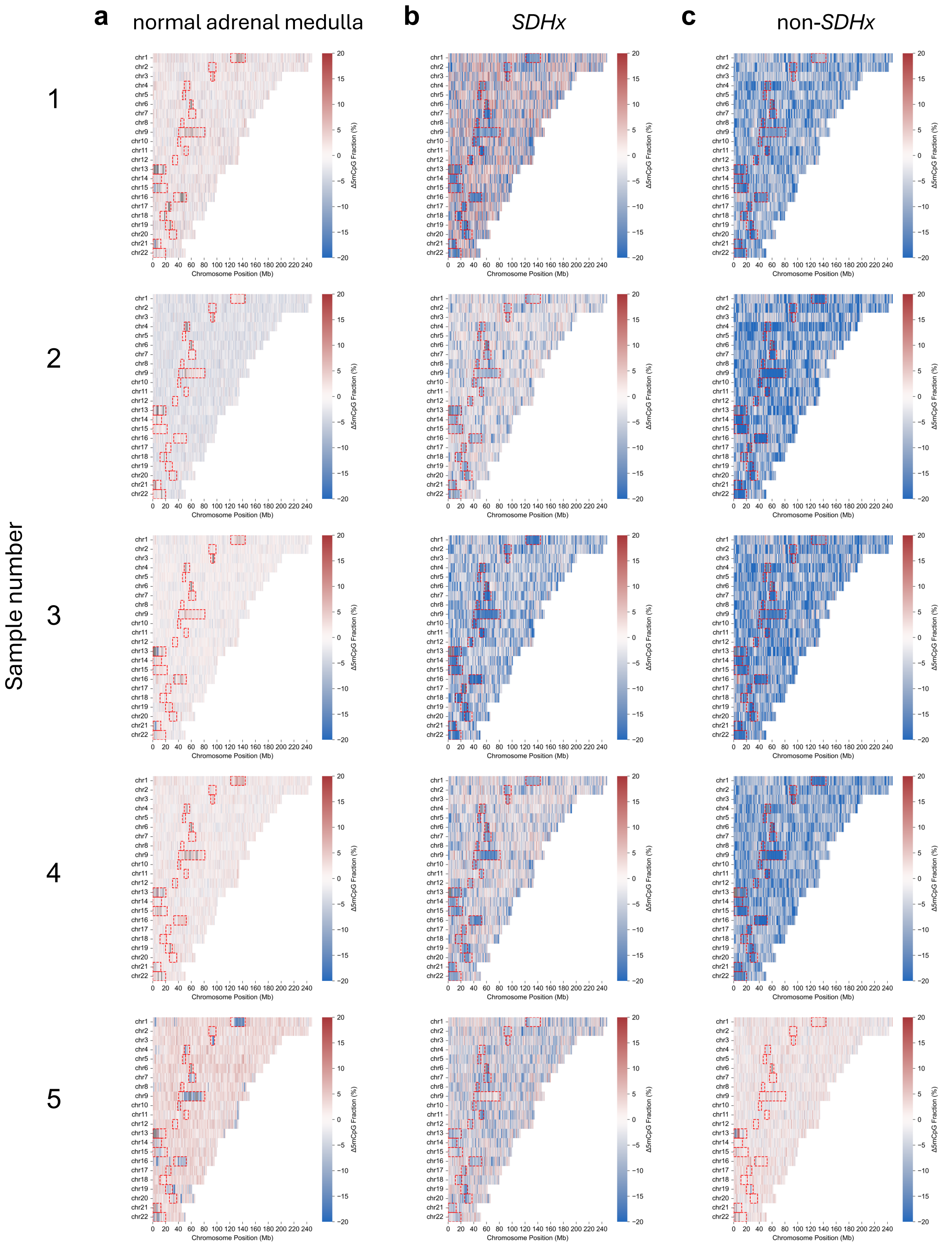


**Supplementary Fig. 2. Differential methylation heatmaps**.

Average methylation in 100-kb windows (sample minus mean normal adrenal medulla cohort). Red indicates hypermethylation; blue indicates hypomethylation relative to normal adrenal medulla. Red boxes: centromeres. (**a**) Normal adrenal medulla. (**b**) *SDHx* tumors. (**c**) Non-*SDHx* tumors.

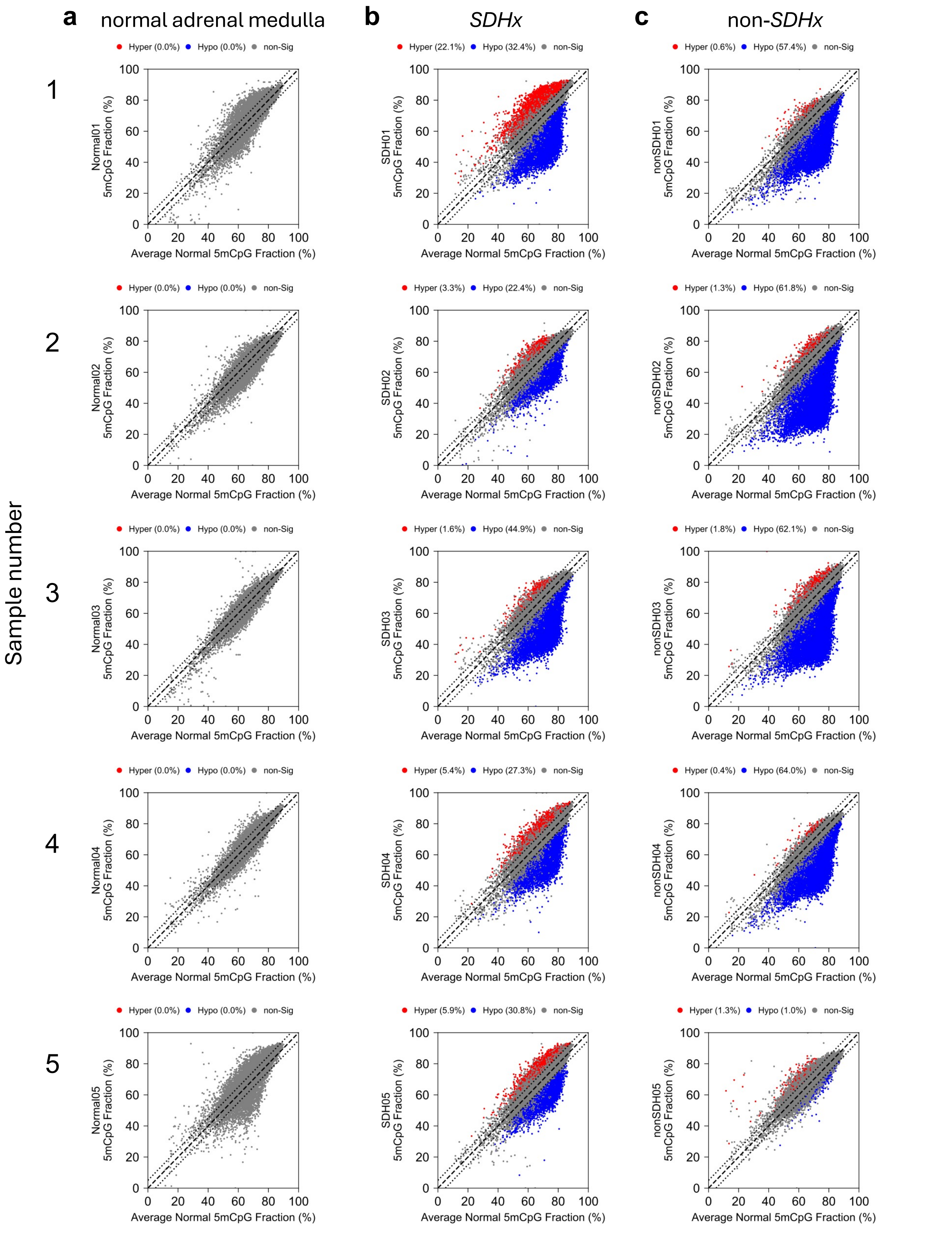


**Supplementary Fig. 3. Scatter plots for autosomal 100-kb windows.** The 22 autosomes were divided into 100-kb windows. The average 5mCpG methylation was computed for each window. Scatterplots comparing methylation in 100-kb windows of individual samples with the normal adrenal medulla cohort average (n = 5). Each dot represents a 100-kb window. Dashed line indicates equal methylation in the sample and the normal cohort average; dotted lines enclose windows differing in methylation by 5% or less. (**a**) Normal adrenal medulla. (**b**) *SDHx* tumors. (**c**) Non-*SDHx* tumors.

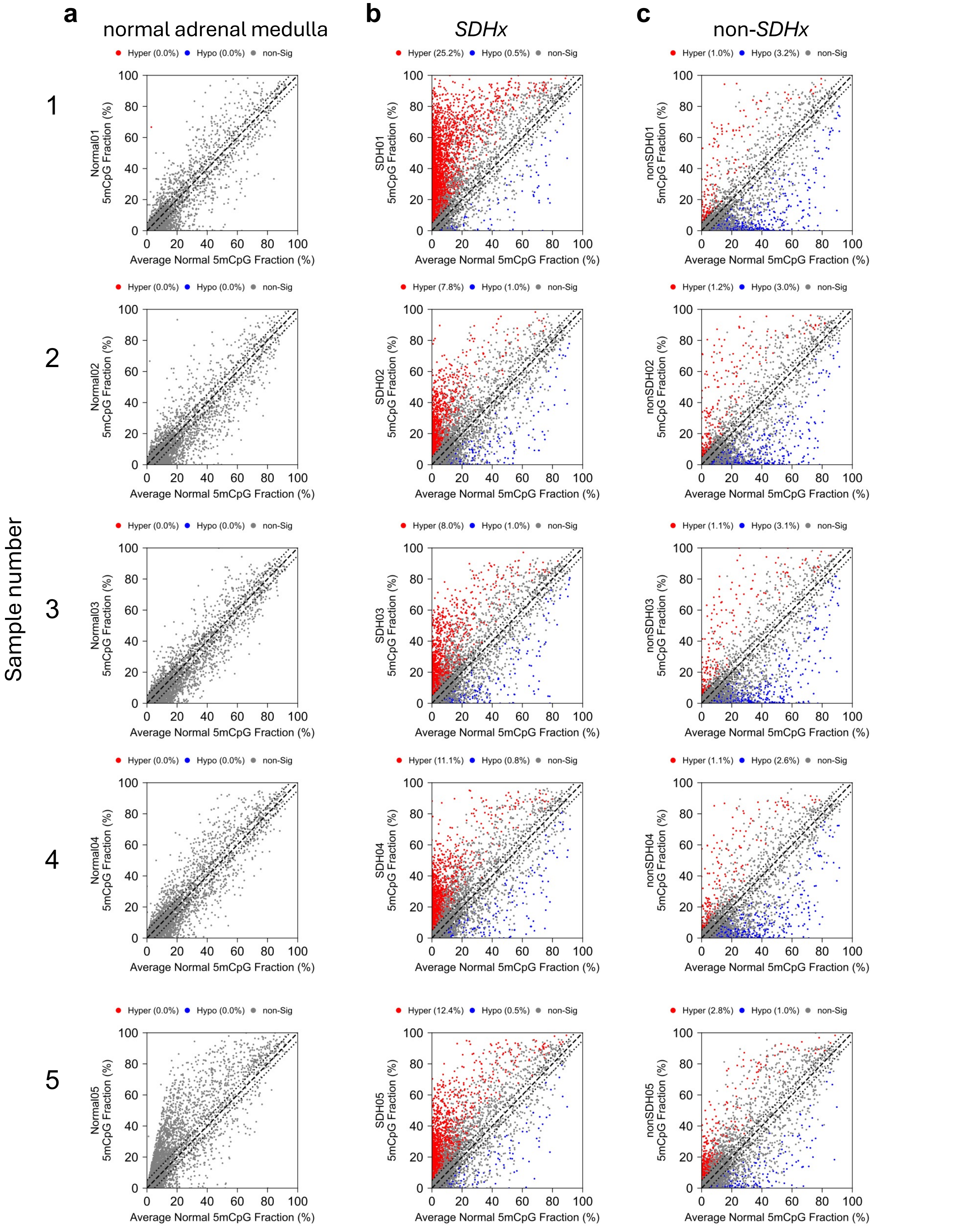


**Supplementary Fig. 4. CpG islands marked with H3K4me3 and/or H3K4me1.** Scatterplots comparing CpG island methylation in individual samples with the normal adrenal medulla cohort average (n = 5). Each dot represents a CpG island. Dashed line indicates equal methylation in the sample and the normal cohort average; dotted lines enclose islands differing in methylation by 5% or less. (**a**) Normal adrenal medulla. (**b**) *SDHx* tumors. (**c**) Non-*SDHx* tumors.

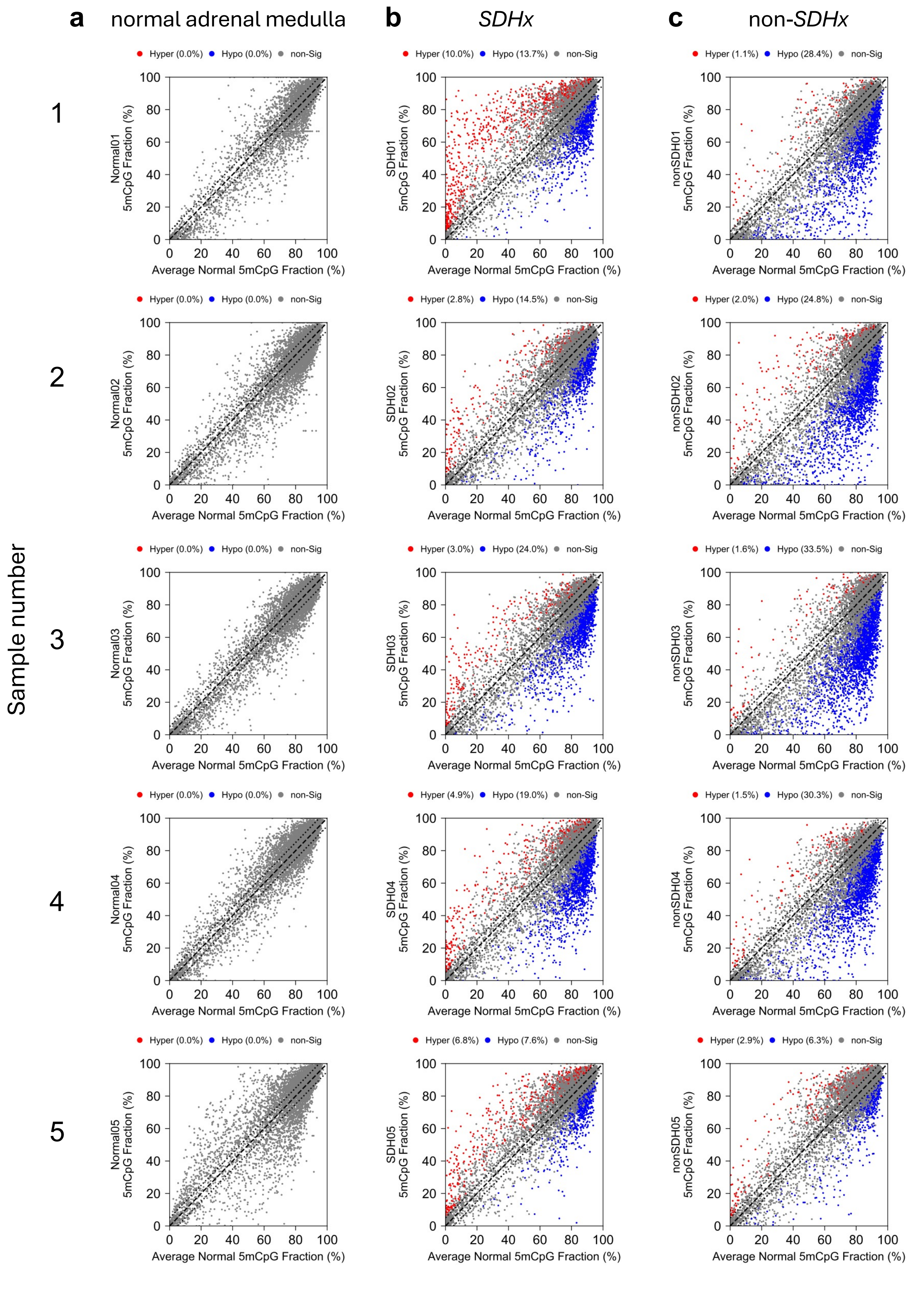


**Supplementary Fig. 5. CpG islands lacking histone marks.**

Scatter plots comparing methylation in CpG islands lacking histone marks (H3K4me1/3, H3K27ac, H3K27me3 or H3K9me3) in individual samples with the normal adrenal medulla cohort average (n = 5). Each dot represents a CpG island. Dashed line indicates equal methylation in the sample and the normal cohort average; dotted lines enclose islands differing in methylation by 5% or less. (**a**) Normal adrenal medulla. (**b**) *SDHx* tumors. (**c**) Non-*SDHx* tumors.

### Supplementary Table 1. List of samples.

| Sample Type | Sample Name | Tissue | Sex | Driver | Average Coverage* | N50 | Average m5CpG (%)** | Tumor Type | Age at surgery |
|---|---|---|---|---|---|---|---|---|---|
| Normal | Normal01 | Adrenal | Male | - | 4.1 | 11345 | 68.8 | - | 51 |
| Normal | Normal02 | Adrenal | Female | - | 4.8 | 6674 | 66.1 | - | 68 |
| Normal | Normal03 | Adrenal | Male | - | 7.8 | 6468 | 68.0 | - | 45 |
| Normal | Normal04 | Adrenal | Male | - | 7.5 | 9525 | 68.6 | - | 62 |
| Normal | Normal05 | Adrenal | Male | - | 5.7 | 14612 | 69.8 | - | 57 |
| *SDHx* | SDH01 | Adrenal | Female | SDHB c.761dup (p.Lys255fs) | 8.5 | 7836 | 65.9 | A-PGL | 50 |
|  | SDH01 | Blood | Female | SDHB c.761dup (p.Lys255fs) | 8.6 | 8252 | 77.7 | - | 50 |
| *SDHx* | SDH02 | Adrenal | Male | SDHD c.337_340del | 7.3 | 6723 | 64.9 | PCC | 47 |
|  | SDH02 | Blood | Male | SDHD c.337_340del | 6.9 | 21637 | 77.5 | - | 47 |
| *SDHx* | SDH03 | Adrenal | Female | SDHB c.137G>A (p.Arg46Gln) | 9.5 | 5572 | 59.0 | A-PGL | 31 |
|  | SDH03 | Blood | Female | SDHB c.137G>A (p.Arg46Gln) | 6.4 | 11435 | 78.4 | - | 31 |
| *SDHx* | SDH04 | Adrenal | Male | SDHD c.242C>T (p.Pro81Leu) | 7.9 | 4211 | 64.6 | CBT | 42 |
|  | SDH04 | Blood | Male | SDHD c.242C>T (p.Pro81Leu) | 6.0 | 10977 | 77.9 | - | 42 |
| *SDHx* | SDH05 | Adrenal | Female | SDHB c.526G>T (p.Glu176Ter) | 10.4 | 5599 | 66.3 | UB-PGL | 64 |
|  | SDH05 | Blood | Female | SDHB c.526G>T (p.Glu176Ter) | 9.0 | 20954 | 78.8 | - | 64 |
| non-*SDHx* | nonSDH01 | Adrenal | Female | RET c.1900T>C (p.Cys634Arg) | 6.8 | 4390 | 57.7 | PCC | 30 |
|  | nonSDH01 | Blood | Female | RET c.1900T>C (p.Cys634Arg) | 7.3 | 20948 | 78.6 | - | 30 |
| non-*SDHx* | nonSDH02 | Adrenal | Female | RET c.1901G>A (p.Cys634Tyr) | 10.6 | 8017 | 54.5 | PCC | 22 |
|  | nonSDH02 | Blood | Female | RET c.1901G>A (p.Cys634Tyr) | 14.0 | 10435 | 78.4 | - | 22 |
| non-*SDHx* | nonSDH03 | Adrenal | Female | RET c.2753T>C (p.Met918Thr) | 8.8 | 9850 | 54.9 | PCC | 48 |
|  | nonSDH03 | Blood | Female | RET c.2753T>C (p.Met918Thr) | 10.7 | 7463 | 78.6 | - | 48 |
| non-*SDHx* | nonSDH04 | Adrenal | Male | NF1 c.3264del (p.Glu1089fs) | 10.2 | 6043 | 56.6 | PCC | 71 |
|  | nonSDH04 | Blood | Male | - | 15.8 | 23145 | 78.0 | - | 71 |
| non-*SDHx* | nonSDH05 | Adrenal | Female | Apparently sporadic (no mutations in PPGL-related genes) | 7.7 | 5925 | 69.0 | HNPGL | 41 |
|  | nonSDH05 | Blood | Female | - | 6.7 | 23507 | 77.5 | - | 41 |

* Measured by average coverage of 100 kb windows in 22 autosomes

** Average % mCpG in the 22 autosomes (Total mCpG / Total All CpG)
